# Homoeologous gene expression and co-expression network analyses and evolutionary inference in allopolyploids

**DOI:** 10.1101/2019.12.16.878900

**Authors:** Guanjing Hu, Corrinne E. Grover, Mark A. Arick, Meiling Liu, Daniel G. Peterson, Jonathan F. Wendel

## Abstract

Polyploidy is a widespread phenomenon throughout eukaryotes. Due to the coexistence of duplicated genomes, polyploids offer unique challenges for estimating gene expression levels, which is essential for understanding the massive and various forms of transcriptomic responses accompanying polyploidy. Although previous studies have explored the bioinformatics of polyploid transcriptomic profiling, the causes and consequences of inaccurate quantification of transcripts from duplicated gene copies have not been addressed. Using transcriptomic data from the cotton genus (*Gossypium*) as an example, we present an analytical workflow to evaluate a variety of bioinformatic method choices at different stages of RNA-seq analysis, from homoeolog expression quantification to downstream analysis used to infer key phenomena of polyploid expression evolution. In general, GSNAP-PolyCat outperforms other quantification pipelines tested, and its derived expression dataset best represents the expected homoeolog expression and co-expression divergence. The performance of co-expression network analysis was less affected by homoeolog quantification than by network construction methods, where weighted networks outperformed binary networks. By examining the extent and consequences of homoeolog read ambiguity, we illuminate the potential artifacts that may affect our understanding of duplicate gene expression, including an over-estimation of homoeolog co-regulation and the incorrect inference of subgenome asymmetry in network topology. Taken together, our work points to a set of reasonable practices that we hope are broadly applicable to the evolutionary exploration of polyploids.

## INTRODUCTION

Comparative transcriptomics has become a widely employed and powerful tool in plant evolutionary biology. Applications are many and diverse, including evolutionary rate estimation [1–3], reconstruction of species relationships [3–5], and the elucidation of co-expression and regulatory changes in gene networks [6, 7]. Next-generation sequencing has facilitated inexpensive and efficient transcriptomic profiling for species whose lack of existing genomic resources would have previously been an obstacle. A landmark example is the recent publication of transcriptomes from more than 1000 species of green plants, which substantially improved available resources and facilitated comparative transcriptomics and phylogenetics among previously underrepresented plants [8](www.onekp.com). This success led to the 10KP project (https://db.cngb.org/10kp/), which aims to sequence 10,000 plant and protist genomes within the next 5 years to further advance our understanding of plant evolution and diversity.

In the context of comparative transcriptomics, polyploid genomes offer unique challenges due to the coexistence of highly similar duplicated genes (homoeologs). Polyploidy in plants is far more prevalent than once thought, acting historically and more recently to shape the genomes of all angiosperms and most other groups of plants [8–11]. One realization that has emerged in the last decade is that polyploidy is accompanied by massive transcriptomic responses, as reviewed [12–14]. These responses are many and varied, including biased homoeolog expression, condition-specific differential homoeolog usage, transgressive expression levels, and expression level dominance. Duplicated gene expression patterns are coordinated in ways that are not fully understood and which depend on myriad factors, including dosage effects, gene balance, interactions among divergent *cis* and *trans*-acting factors, and various topological aspects of gene networks [6,15–18].

Research on polyploid transcriptomes is divided into two broad categories with respect to the treatment of homoeologs: those that evaluate individual homoeolog expression separately and those that evaluate the aggregate expression of homoeologs. The ability to consider homoeologs separately depends largely upon the mode of origin (autopolyploid vs. allopolyploid) and the extent of sequence divergence between homoeologs, as well as the genomic resources available. Distinguishing individual homoeolog expression levels is difficult when sequence divergence between homoeologs is too low, as often is the case with allopolyploids formed from recently diverged diploid parents, or in evolutionarily young autopolyploids. When a reference genome or transcriptome is available for a polyploid, quantitation of individual homoeolog expression levels is possible if sequence divergence is sufficiently high, and aggregated expression can be derived from the summation of each homoeolog set. In many cases, reference genomes may only be available for one or more model diploids. These diploid genomes can be useful in analyses of duplicate gene expression in polyploids, but they require additional steps to characterize and partition homoeolog-specific reads. Regardless of the ploidy level of the reference genome, short RNA-seq reads may be difficult to explicitly map to individual homoeologs due to their near-duplicate nature (i.e., multi-mapped reads). That is, only a certain proportion of reads (related to divergence between homoeologous genomes) will contain homoeolog distinguishing variants (e.g. SNPs). Only those reads that can be unambiguously assigned to specific homoeologs can be utilized for homoeolog transcript counting (Figure 1A).

**Figure 1.**
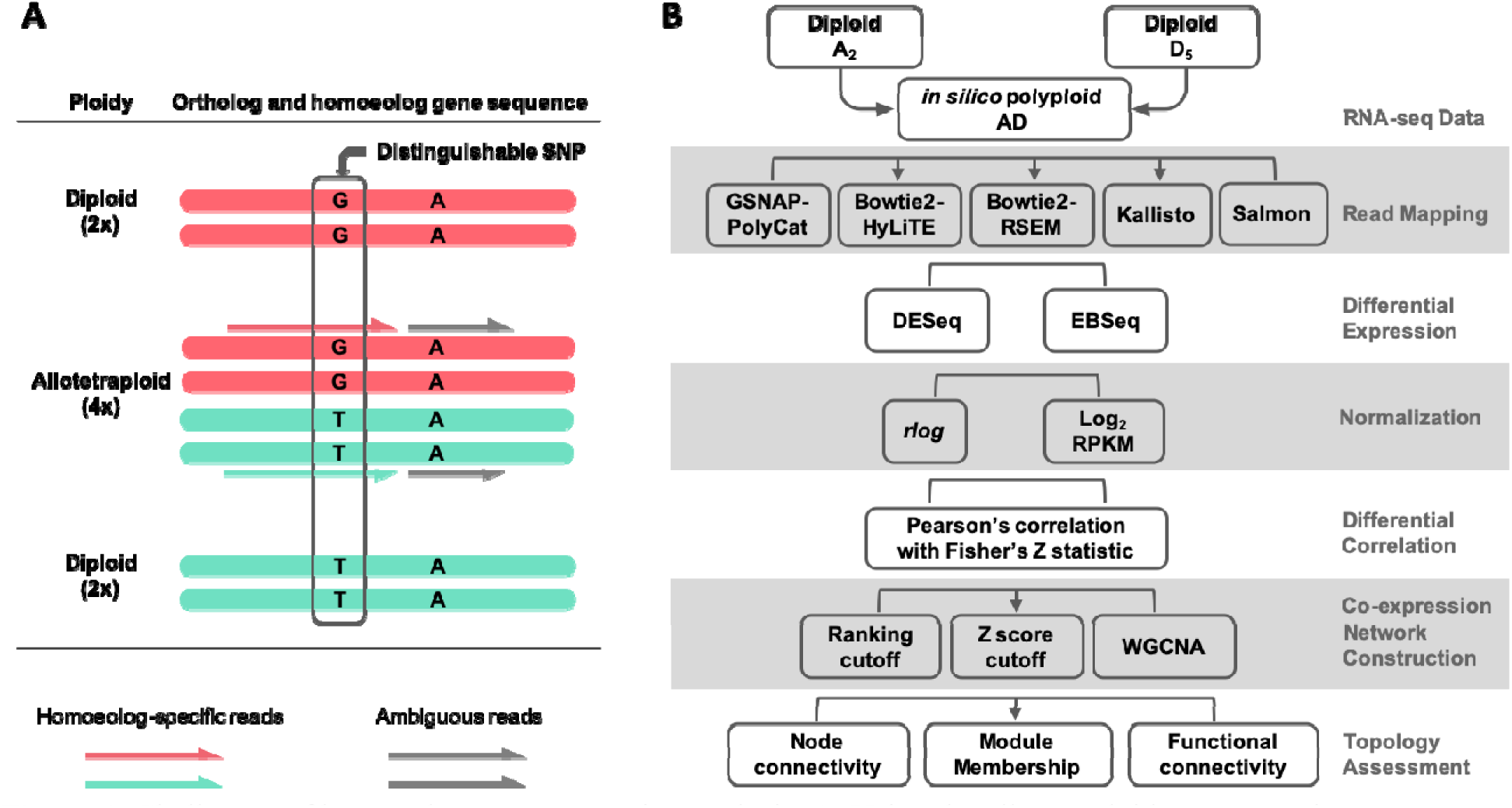
Challenges of homoeolog gene expression analysis. **A**. Using the allotetraploid cotton species as an example, only a small portion of RNA-seq reads contain diagnostic SNPs (i.e., homoeolog-specific reads) reflecting the parental origin of homoeologous genes. **B**. An analytic workflow of RNA-seq analysis was applied to evaluate the use of homoeolog-specific reads to study duplicated gene expression and co-expression networks. A ground-truth, *in silico* dataset of allopolyploid cotton (AD) was generated from the parental diploid cotton A_2_ and D_5_ reads, which was analyzed using a variety of method choices.

As previously noted by Ilut et al. [19], the issue of ambiguous read mapping is prevalent in plants due to their natural genomic redundancy, and is even more so for recent polyploids. Many intrinsic and extrinsic factors affect the ability to partition homoeolog expression, including: (1) divergence between subgenomes, in terms of frequency and distribution of SNPs; (2) the sequencing strategy (e.g. read length, and paired- vs. single-ended reads) for generating RNA-seq data; (3) the quality of reference genome(s); and (4) the bioinformatic tools used for partitioning and/or quantifying homoeolog-specific reads, including methods for allocating ambiguous reads in general (such as RSEM [20] and Salmon [21]) and those specifically developed for polyploid systems (i.e., PolyCat [22], PolyDog [23], HyLiTE [24], HANDS [25] and HAND2 [26]).

Given these complexities inherent in working with polyploid transcriptome data, the question arises as to how these factors affect our ability to derive accurate polyploid gene expression profiles. That is, how do the many issues noted above affect read assignment and our inferences of gene expression and co-expression characteristics? Here we explore the causes and consequences of read ambiguity in homoeologous differential expression and co-expression networks using transcriptome data from the cotton genus (*Gossypium*) as an example (Figure 1B). Tetraploid cotton (represented here by *G. hirsutum*; AD_1_) originated from an allopolyploidization event between an A-genome (*G. herbaceum*- or *G. arboreum*-like) and a D-genome (*G. raimondii*-like) diploid species circa 1 to 2 million years ago (reviewed in [27]). Because there is no gold standard for true expression levels of At and Dt (t denotes subgenome) homoeologs in the polyploid AD_1_ transcriptomes, we generated *in silico* allopolyploid datasets (AD) by combining reads from the A- and D-genome diploid transcriptomes (see methods). This approach allowed us to evaluate accuracy of “homoeolog” expression against the actual diploids reads used for generating *in silico* dataset. Methodologically, we first evaluated a variety of bioinformatic method choices at different stages of RNA-seq data analysis, with the aim of generating insight into best practices that may be broadly applicable to other polyploid systems.

## METHODS

All codes used in this study are available in Github https://github.com/Wendellab/homoeologGeneExpression-Coexpression.

### Data availability and preparation

For generating *in silico* allopolyploid cotton (AD) datasets, matched RNA-seq data of the model diploid progenitors, i.e., *G. arboreum* (A_2_) and *G. raimondii* (D_5_), were obtained, each comprising 33 RNA-seq libraries under 12 sample conditions (Table S1). The seed dataset under NCBI BioProject PRJNA179447 consists of 11 libraries (4 seed developmental stages each with 2-3 biological replicates) for each diploid with 100 bp single-end reads and an average of 14.8 million reads per library. The flowering dataset under NCBI BioProject PRJNA529417 consists of 22 libraries (8 tissues each with 2 to 3 biological replicates) for each diploid with 150 bp paired-end reads and an average of 13.8 million read pairs per library. Following adaptor and quality trimming via Sickle [v1.33] [28], the matched A_2_ and D_5_ libraries (at each condition and replicate) were adjusted to contain equivalent number of filtered reads and subsequently combined to generate the corresponding *in silico* allopolyploid (AD) datasets. For each pair of AD homoeologous genes, the gene regions that should be unambiguously assigned to subgenome (i.e. effective region), given the distinguishable SNP distribution between homoeologs and the specific sequencing strategy involved, were detected using a custom script “detectEffectiveRegion.r”. The proportion of each gene sequence that belongs to an effective region was calculated as *%Eflen*. We next introduced a metric of *Ambiguity* for each pair of homoeologous genes as calculated by 1-*%Eflen*, because *%Eflen* is inversely correlated with the number of ambiguous reads that cannot be assigned via direct SNP evidence.

**Table 1.** Nine classes of differential correlation (DC) changes.

| Diploid correlation | Polyploid correlation | Description of DC pattern |
| --- | --- | --- |
| + | + | Both positive but different in $r$-value |
| + | 0 | Loss of positive correlation |
| + | - | Inversion from positive to negative correlation |
| 0 | + | Gain of positive correlation |
| 0 | 0 | Neither significant but different in $r$-value |
| 0 | - | Gain of negative correlation |
| - | + | Inversion from negative to positive correlation |
| - | 0 | Loss of negative correlation |
| - | - | Both negative but different in $r$-value |

### RNA-seq read mapping and homoeolog-specific read partitioning

The following five pipelines were each independently applied to the diploid and AD polyploid datasets.

#### GSNAP-PolyCat

This pipeline utilizes the SNP-tolerant capabilities of GSNAP [v2016-08-16] [29] to map polyploid reads to a single diploid progenitor genome (here, *G. raimondii*; [30]). The SNP-tolerant feature of GSNAP permits equivocal mapping of both A- and D-diploid derived reads based on *a priori* SNP information. Here, we used a previously generated genome-diagnostic SNP-index [22] for mapping. The resulting alignments were sorted using samtools [31] and subsequently partitioned into homoeolog-specific reads using PolyCat [v1.3] [22]. Read counts were tabulated using HTSeq [v0.9.1] [32].

#### HyLiTE

This software automates the process of read mapping, SNP detection, and read count partitioning in a single step [24]. Briefly, HyLiTE [v.2.0.1] employs Bowtie2 [v2.3.1] [33] to map both diploid and polyploid reads to the reference gene models (here *G. raimondii*; (Paterson *et al.*, 2012)), and sorts homoeologous reads based on the SNPs detected from mapping the diploid reads. Homoeolog-specific read count tables are automatically generated in the last step.

#### RSEM

While not specifically developed for polyploids, RSEM [v1.3.0] [20] and the following programs (i.e., Salmon [v0.9.1] [21] and Kallisto [v0.44.0] [34]) were developed to address the general issues of ambiguously mapped reads while also increasing mapping speed. RSEM automates read alignment to a set of reference transcripts using Bowtie2 and subsequently estimates feature counts using the EM algorithm, both at the gene and isoform level. As the presence of homoeologs is bioinformatically similar to presence of alleles of isoforms, RSEM may be suitable for disentangling homoeologous reads and estimating homoeolog abundance. For RSEM, we approximated the polyploid reference transcriptome by combining the *G. raimondii* transcriptome and the predicted *G. arboreum* (A_2_) transcripts based on the same SNP index used by GSNAP-PolyCat. That is, the *G. arboreum* transcripts here are simply the *G. raimondii* transcripts with homoeologous SNP sites replaced with *G. arboreum-*specific SNPs.

#### Kallisto

This method belongs to a class of read aligners known as “pseudoaligners”, which leverage kmer information to detect the transcripts that could have generated a given read without specifically aligning the read [34]. Kallisto, like other pseudoaligners, generates a de Bruijn graph of the kmers present in a transcriptome to quickly assign reads based on intersecting read and transcriptome kmer metrics. Kallisto was run under default parameters using the above generated polyploid reference transcriptome.

#### Salmon

This method employs a light-weight, quasi-mapping strategy [35] similar to Kallisto and a two-phase estimation of expression. This two-phase estimation uses two forms of Bayesian inference (Foulds et al. 2013; Do and Batzoglou 2008) to first estimate and then subsequently refine transcript-level abundances [21]. Using this method, Salmon is able to estimate abundance uncertainty due to ambiguously mapped reads, which are common with homoeologs. Salmon was also run with default parameters using the above generated polyploid reference transcriptome and the option “keepDuplicates” for indexing the transcriptome. Estimated transcript abundance is automatically returned by the program.

### Performance evaluation of estimating homoeolog expression

For each set of bioinformatically partitioned reads, multiple measures of performance were conducted. Because the true assignment of each *in silico* polyploid (AD) read is known and originates from only two sources (A_2_ and D_5_), assessing homoeolog assignment becomes a binary classification problem. For example, when classifying A_2_-derived reads from the synthetic polyploid transcriptome reads (i.e., At reads), the read could either be correctly assigned to At (true positive; *TP*) or incorrectly assigned to Dt (false negative; *FN*). The same applies to D_5_-derived reads; a D_5_-derived reads assigned to Dt is a *TP*, whereas the assignment to At is a *FN*.

The prediction results of the binary classification can be arrayed as a 2×2 confusion matrix, which summarizes the numbers of true/false positives/negatives (*TP*, *FP*, *TN* and *FN*) that can be evaluated using information retrieval statistics [36], such as Precision/Recall [37] and the Matthews correlation coefficient (*MCC*) [38]. The general formulas of these statistics are as follows: 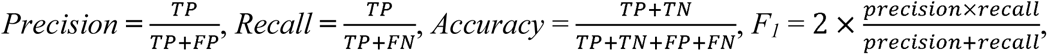 and 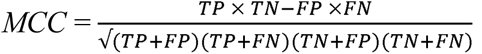. Here we report both the *F_1_* and *MCC* scores, which provide a generalized measure of accuracy; however, we note that *MCC* may be preferred because it accounts for more of the confusion matrix and is more balanced with respect to classes of very different sizes [39].

We also note that the results of binary classification measures for GSNAP-PolyCat and HyLiTE are somewhat misrepresentative of those pipelines. Because GSNAP-PolyCat and HyLiTE discard reads with no diagnostic SNPs, the number of *TPs* and *FNs* will be distorted for these pipelines, i.e., reduced and increased, respectively. In contrast, the remaining pipelines (i.e., RSEM, Salmon, and Kallisto) use statistical inference to completely assign all reads to homoeologs. We therefore define two additional measures that reflect these differences, *Efficiency* and *Discrepancy*. Here, the measure of *Efficiency* is simply the number of reads assigned to a homoeolog class (regardless of accuracy) divided by the total number of reads. The overall difference between the obtained read count and expected true read count for each class were measured by their 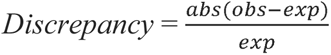.

### Gene expression analysis

Two methods of differential expression (DE) were used to analyze homoeologs expression, i.e., DESeq2 [40] and EBSeq [41]. DESeq2 takes a classical hypothesis testing approach to report nominal *p*-values, whereas EBSeq accommodates the uncertainty inherent in isoforms (here, homoeologs) using a Bayesian framework to return posterior probabilities for differential expression. A false discovery rate α < 0.05 was required to determine significant DE changes, which was applied to the adjusted *p*-values of DESeq2 [42] and the posterior probability (=1-α) of EBSeq.

Because these *in silico* polyploid data were derived from combining diploid libraries, the null hypothesis is that DE between inferred homoeologs should match the DE observed between the diploid libraries for those genes. We again treat this as a binary classification problem, marking each gene as DE or non-DE and comparing the observed number of DE genes in the polyploid libraries with the expected number derived from the diploid data. The same statistical measures of performance (i.e., *Precision*, *Recall*, *Accuracy*, *F_1_,* and *MCC*) were calculated for each pipeline, as described above. The receiver operating characteristic (ROC) curve and the area under the ROC curve (AUC) were calculated for each and visualized using the R package ROCR [43]. AUC scores reflect the probability that a random classification is correct, ranging from 0.0 to 1.0 [44, 45].

### Gene expression correlation analysis

Differential correlation (DC) analysis is commonly used to evaluate coordinated changes in gene expression, either independent of, or in the context of, co-expression network analyses. Both DC and network analyses require some form of variance-stabilizing transformation of the raw data. Several methods of normalization exist [46, 47], which have their own advantages and nuances. Here two common methods were tested, i.e., RPKM followed by log_2_ transformation and regularized logarithm (*rlog*) transformation as implemented in DESeq2.

Using the R package DGCA [48], Pearson correlation coefficients (*r*) and their corresponding *p* values were calculated for each pair of genes across multiple samples, which were subsequently classified as having a significant (*p* < 0.05) positive correlation (+), a significant negative correlation (-), or not significantly different from zero (0). Fisher’s *Z*-test [49] was used to identify significant correlation changes between the homoeologous and the diploid (expected) *r* values.

Given that each condition (i.e., diploid or polyploid) exhibits three possible correlation conditions (+, -, or 0), there are nine possible categories to describe the pattern of DC (Table 1). Among those, 0/0, +/+ and -/- indicate significant changes in *r* values while the inference of correlation condition remains unchanged, and the other classes indicate that the read partitioned AD dataset mis-identities the true condition of gene-to-gene correlations. We assessed enrichment of each class for each pipeline using a one-sided Fisher’s exact test (*p* < 0.05).

Finally, as previously described [15], we compiled a list of genes that are overrepresented with the gene-to-gene paired DC relationships (see above) to identify differentially co-expressed genes (DC genes). Briefly, the probability *p* of any pair of genes exhibiting a DC relationship is defined as the percent of DC pairs detected among all possible gene pairs. For a gene observed in *k* DC pairs among all possible pairs *n*, the probability of a “differential co-expression gene” follows the binomial distribution model:

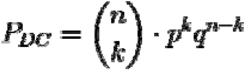

*P_DC_* was corrected by the BH method [42] and a cutoff of 0.05 was used for identifying DC genes.

### Co-expression network construction

Co-expression networks are a multidimensional representation of the expression relationships among genes. Accordingly, construction of co-expression networks use similarity scores from the pairwise gene expression profiles to generate an adjacency matrix which reflects connections between genes (as nodes) in network [50]. Here, we used the Pearson correlation coefficients to calculate the matrix of similarity scores. Derived from this correlation matrix, the adjacency matrix was used as the basis for a series of binary and weighted gene co-expression networks. which were generated for both the log_2_RPKM- and *rlog*-transformed read count tables from each expression estimation pipeline.

For constructing binary networks, a hard threshold was applied to similarity scores to determine whether a pair of genes should be connected in the network, resulting an adjacency matrix containing only 0 and 1 values. Two types of hard thresholds were tested, specifically rank-based and Fisher’s *Z*-statistics [49] based thresholds. A set of rank-based cutoffs (5%, 1%, 0.5%, and 0.1%) were applied to these similarity scores in order to select the top ranked connections as possible edges. Following Fisher’s *Z* transformations to convert each Pearson correlation coefficient to a *Z*-score, a set of cutoffs (i.e., 1.5, 2.0, 2.5, and 3.0) were used to retain correlations with *Z*-score above the cutoff value as edges. The performance of network construction was evaluated as a binary classification problem; that is, because we expected to see the edges inferred from the expected expression (diploid) retained in the polyploid network, we were able to create a confusion matrix from the presence or absence of edges compared to what was expected. The edge classification was again evaluated with a ROC curve using the R package ROCR [43]. Due to the large gene number in the network (> 60,000 genes), a 10% random sampling of genes was used for computation with 10 iterations.

While binary networks have their own utility, weighted co-expression networks are frequently used for reasons enumerated elsewhere [51], including the ability to quantify network connections. Weighted networks use soft thresholding to assign connection strengths to gene pairs, thereby allowing the adjacency matrix to present network connections quantitatively. Using the R package WGCNA [52, 53], a set of soft thresholds (1, 12, 24) were applied for automatic network construction with the *blockwiseModules* function and the following parameters: corType = “pearson”, networkType = “signed”, TOMType = “signed”, minModuleSize = 100. The performance of each polyploid network construction was evaluated against the reference network generated using the diploid data. Preservation of the reference network modules by AD dataset was calculated using the WGCNA function *modulePreservation* with 200 permutations. In general, modules with the derived preservation score *Z_summary_* > 10 are interpreted as strong preservation.

### Network topology measures and functional connectivity assessment

Node connectivity and functional connectivity are two metrics that may provide insight into the importance and/or function of a given gene in a network. Node connectivity (*k*) measures the connectivity of any given node in the network, either by counting the number of connected edges (for a binary network) or summing the connected edge weights (for a weighted network).

Functional connectivity (*FC*) uses the ‘Guilt-by-Association’ principle to measure network quality under the assumption that genes with similar functions should be connected in a well-constructed network. A neighbor voting algorithm from the R package EGAD [54] was used to classify genes into functional groups based on the functionality of their connected genes (i.e., their neighborhood). This package uses the the known functional labels of genes (e.g. GO and KEGG annotations) and the voting algorithm as a binary classifier to return true or false predictions for those functional labels; the performance of the neighbor voting functional assignment can then be assessed by an ROC curve. The derived AUC characterizes the degree to which an input network topology can predict the gene membership of a functional category, which intuitively corresponds to the assessment of functional connectivity. GO and KEGG terms were extracted from the v2.1 annotation of *Gossypium raimondii* reference genome downloaded from Phytozome (www.phytozome.net).

## RESULTS

### Subgenome divergence and homoeolog read ambiguity: the problem

As mentioned in the introduction, the issue of ambiguous read mapping is prevalent in polyploids and in plants in general because of means other than polyploidy that generate paralogs. Accurately partitioning polyploid reads is bioinformatically challenging (Figure 1A), and the consequences of inaccurate partitioning are unknown. The proportion of ambiguous reads is dependent both on subgenome divergence and the sequencing strategy, and the subsequent treatment (i.e., removal or statistical assignment) can affect the outcome of downstream analyses. Here we evaluated the performance of five different pipelines in assigning reads to polyploid genomes and the effects of their treatment of ambiguous reads on downstream analyses of duplicated gene expression (Figure 1B). Accordingly, we introduced the metric of *Ambiguity* for each pair of homoeologous genes, which corresponds to the percentage of a gene region that cannot be distinguished between homoeologs (see Methods). Ideally, if the homoeologous sequences were sufficiently divergent and the sequencing reads were long enough to consistently contain homoeolog distinguishable variants (e.g. SNPs), all reads could be assigned with zero *Ambiguity*; however, these conditions are rarely met by existing data.

In tetraploid *Gossypium*, where the average sequence divergence (in coding regions) between homoeologs is approximately 1.5% [22], only 5% of homoeologous gene pairs can be unambiguously mapped (*Ambiguity* = 0) by 50 bp RNA-seq reads, whereas over 90% can be unambiguously mapped by 300 bp reads (Figure 2A). In the following analysis, we binned homoeologous gene pairs into five increasing levels of *Ambiguity*, i.e., (0), (0-0.05), (0.05-0.1), (0.1-0.2), and (0.2-1.0). These bins were next used to relate the performance of read assignment and other duplicated gene expression patterns to the level of read ambiguity (Figure 2).

**Figure 2.**
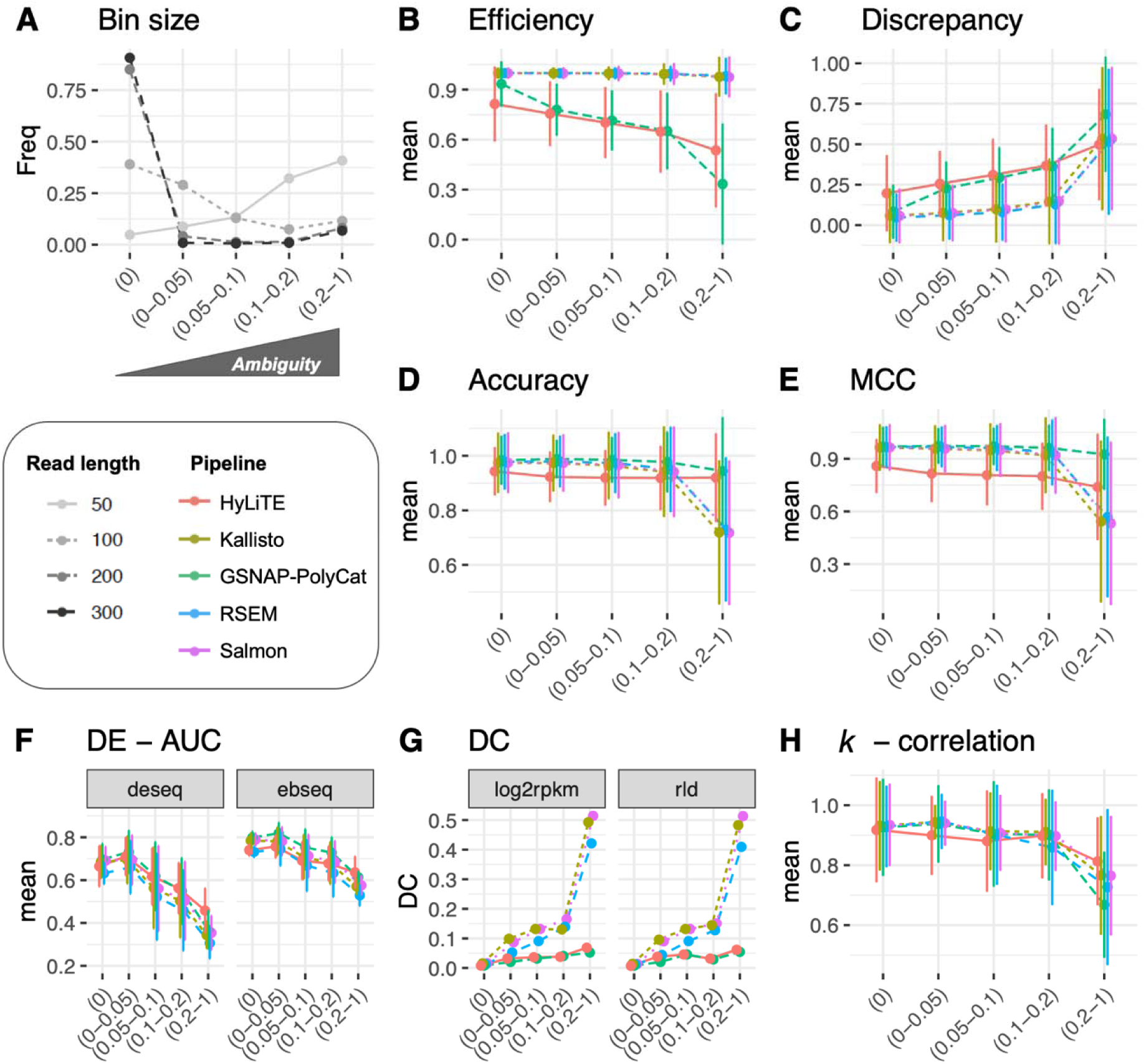
Homoeologous read ambiguity and consequences. **A**. Given the specific sequencing read length (i.e. 50, 100, 200 and 300 bp), the homoeologous gene pairs from *Gossypium* were binned by *Ambiguity* into five groups: (0), (0-0.05), (0.05-0.1), (0.1-0.2), and (0.2-1.0), the first of which indicates complete read assignment via SNP differentiation. The y-axis refers to the bin size of each gene group. These *Ambiguity* bins were used to relate the performance of read assignment (**B**-**E**), differential expression (**F**), differential correlation (**G**), and the analysis of node connectivity *k* (**H**). Error bars represent the standard deviation.

### Artificial allopolyploid datasets permit assessment of fidelity in homoeologous read assignment

We generated *in silico* allotetraploid (AD) datasets for multiple sample conditions (tissues, developmental timepoints, etc.; Supplementary Table S1) as a ground truth reference. For these, we combined equal amounts of reads from two diploids, *G. arboreum* (A_2_) and *G. raimondii* (D_5_), which represent the model diploid progenitors for a clade of naturally occurring polyploids in *Gossypium*. As these datasets are diploid-derived, the amount of A- and D-derived reads in the AD datasets is known and the ability of each pipeline to accurately reconstruct this becomes testable.

Because the five pipelines differ in how they treat ambiguous reads, either discarding them (GSNAP-PolyCat and HyLiTE) or statistically partitioning them (RSEM, Salmon and Kallisto), we first evaluated the *Efficiency* and *Discrepancy* of read assignment. *Efficiency* simply measures the percentage of reads assigned, considering all the reads versus those partitioned into each subgenome. As shown in Table 2, RSEM, Salmon, and Kallisto all achieved 100% read assignment due to their underlying statistical inference of origin for ambiguous reads; however, they tend to slightly overestimate the number of At reads. Since GSNAP-PolyCat and HyLiTE discard ambiguous reads, their *Efficiency* negatively correlates with *Ambiguity,* as expected (Figure 2B), with only 87.7% and 82.2% of total reads partitioned into subgenome (Table 2). In contrast to RSEM, Salmon, and Kallisto, there appears to be a reference bias in both GSNAP-PolyCat and HyLiTE that leads to more reads being characterized as D-derived; this bias is most significant for HyLiTE (Table 2; At 78.5% vs. Dt 85.8%, Student’s T test *p* < 0.05). We also evaluated the *Discrepancy* for each pipeline, which measures the absolute difference between the obtained homoeolog read counts and the expected counts; this measure is affected by both the assignment *Efficiency* and binary classification measures (see methods). Due to the 100% *Efficiency* guaranteed by the algorithms of RSEM, Salmon and Kallisto, these pipelines exhibit the lowest *Discrepancy* (5.1%; Table 2), while the highest *Discrepancy* was found in HyLiTE (18.1%), followed by 12.7% in GSNAP-PolyCat; in the latter two, the *Discrepancy* reflects both assignment error and discarded reads. In general, *Discrepancy* from the actual read numbers increases as the level of *Ambiguity* increases (Figure 2C), as expected.

**Table 2.** Overall and subgenome assessment of homoeolog expression estimation. The best performance for each metric is marked in **bold** text.

|  | GSNAP-PolyCat | HyLiTE | RSEM | Salmon | Kallisto |
| --- | --- | --- | --- | --- | --- |
| **Efficiency** | 87.7±1.5% | 82.2±0.7% | **100.0%** | **100.0%** | **100.0%** |
| At | 86.7±1.6% | 78.5±1.0% | 101.0±0.6% | 101.6±0.5% | 101.5±0.6% |
| Dt | 88.6±1.6% | 85.8±0.6% | 99.1±0.6% | 98.5±0.5% | 98.6±0.6% |
| **Discrepancy** | 12.7±1.5% | 18.1±0.8% | **5.1±0.6%** | **5.1±0.6%** | **5.1%±0.6%** |
| At | 13.4±1.6% | 21.±1.06% | 5.4±0.6% | 5.2±0.6% | 5.2±0.6% |
| Dt | 12.1±1.5% | 14.8±0.6% | 4.8±0.6% | 5.1±0.6% | 5.0±0.6% |
| **Precision** | - | - | - | - | - |
| At | **98.4±0.1%** | 92.4±0.3% | 95.1±0.2% | 94.3±0.2% | 94.4±0.2% |
| Dt | **97.7±0.5%** | 91.2±0.6% | 97.0±0.5% | 96.5±0.4% | 96.4±0.4% |
| **Recall** | - | - | - | - | - |
| At | **97.7±0.5%** | 90.6±0.6% | 96.5±0.5% | 96.4±0.4% | 96.4±0.4% |
| Dt | **98.1±0.1%** | 94.0±0.3% | 96.2±0.2% | 95.4±0.2% | 95.4±0.2% |
| **F<sub>1</sub> score** | - | - | - | - | - |
| At | **98.7±0.2%** | 92.2±0.4% | 97.2±0.3% | 96.8±0.2% | 96.9±0.2% |
| Dt | **98.7±0.2%** | 93.1±0.4% | 97.2±0.3% | 96.7±0.2% | 96.7±0.2% |
| **Accuracy\*** | **98.2±0.2%** | 93.7±0.7% | 96.3±0.3% | 95.7±0.2% | 95.8±0.2% |
| **MCC\*** | **96.8±0.5%** | 84.5±0.7% | 95.5±0.5% | 94.7±0.4% | 94.8±0.4% |

\* Same values for At and Dt reads.

While *Efficiency* and *Discrepancy* provide generic measures of read partitioning based on the numbers expected, they do not account for whether each read is accurately assigned. Therefore, the results of each pipeline were arrayed in a 2×2 confusion matrix (i.e., true positive, false negative, etc; [36]) and the performance of the pipeline was evaluated using the information retrieval metrics of *Precision*, *Recall*, *Accuracy*, *F_1_* score, and *MCC*. In the context of information retrieval (as implemented here), *Precision* measures how many of the reads assigned to a given subgenome (A or D) were correctly identified, *Recall* measures how many of each subgenome were retrieved from the mixed population (relative to expectations), and *Accuracy* measures how well each pipeline correctly identifies one subgenome while excluding the other; the measures *F_1_* and *MCC* account for more of the confusion matrix and attempt to generalize the results into a single score of performance (see methods for details). The results in Table 2 show that that GSNAP-PolyCat generally performed better in all information retrieval metrics, meaning that it recovered more relevant reads for each subgenome while excluding reads from the other subgenome. The three generic, alignment-based approaches (i.e., RSEM, Salmon, and Kallisto) showed comparable performance to each other, with only a slight reduction in all scores relative to GSNAP-PolyCat. Only HyLiTE stands out as performing relatively poor compared to the other pipelines; however, it is noteworthy that the other four pipelines all utilized the same SNP information derived from rich genomic resequencing data [23], whereas HyLiTE conducted on-the-fly SNP calling from the input parental diploid RNA-seq datasets. This most likely explains the relatively poor performance of HyLiTE, as tested here. Interestingly, as shown in Figures 2D and 2E, GSNAP-PolyCat and HyLiTE both exhibit relatively consistent performance across *Ambiguity* bins, indicating that their accuracy (as measured by *Accuracy* and *MCC*) is largely static, irrespective of homoeolog divergence. RSEM, Salmon, and Kallisto, however, perform nearly as well as GSNAP-PolyCat when the expected amount of homoeologous ambiguity is low, but quickly descend when *Ambiguity* goes above 20% (Figures 2D and 2E).

### The inference of homoeolog expression divergence is affected by the choice of expression estimating pipeline

Expression divergence of homoeologs, both relative to one another and to their progenitor genomes, is a major component of polyploid research. Allopolyploidy reunites formerly diverged genes (and their regulatory context) into a common nucleus while simultaneously generating massive redundancy. Consequently, observed transcriptomic changes are myriad (reviewed in [13]), and include homoeolog expression bias (reviewed by [12, 13]) and functional divergence [55–58]. Since our ability to accurately describe expression changes depends upon our ability to accurately represent expression, we evaluated the extent to which each pipeline accurately represented differential expression (DE) between homoeologs. That is, the homoeolog DE results derived from each pipeline inferred AD dataset were compared to the expected DE results between the diploid orthologs from which the AD dataset was derived. While many methods exist for comparing DE among samples, we selected two of the most popular methods, namely DESeq2 and EBSeq, to compare both stringency and accuracy in general and in the context of the different pipelines.

Overall, DESeq2 detected an average of 18% more significant changes in expression than EBseq (Supplementary Table S2; paired Student’s T test *P* < 0.05), suggesting that by default the latter is more stringent. Across the twelve sample conditions, homoeolog expression divergence was detected from between 5% and 44% of the 37,223 homoeologous gene pairs, without significant differences between the observed and expected results (Supplementary Table S2; paired Student’s T test *p* = 0.26). As shown in Figure 3, a relatively high level of DE *Accuracy* (above 80%) was consistently inferred. Regardless of which DE method was used, the expression datasets generated by GSNAP-PolyCat outperformed those by other pipelines (Salmon/Kalisto > HyLiTE/RSEM) in identifying the true expression divergence between homoeologs. While DESeq2 appeared to perform better than EBseq according to the measures of *Precision*, *Recall, F_1_* and *MCC*, the AUC scores suggested that EBseq is more robust than DESeq2 to separate binary classes (Figure 3), particularly for genes exhibiting high *Ambiguity* (Figure 2F). For both methods, their AUCs were negatively correlated with *Ambiguity*, reflecting the strong dependence of DE analysis on the extent of homoeolog sequence divergence (Figure 2F).

**Figure 3.**
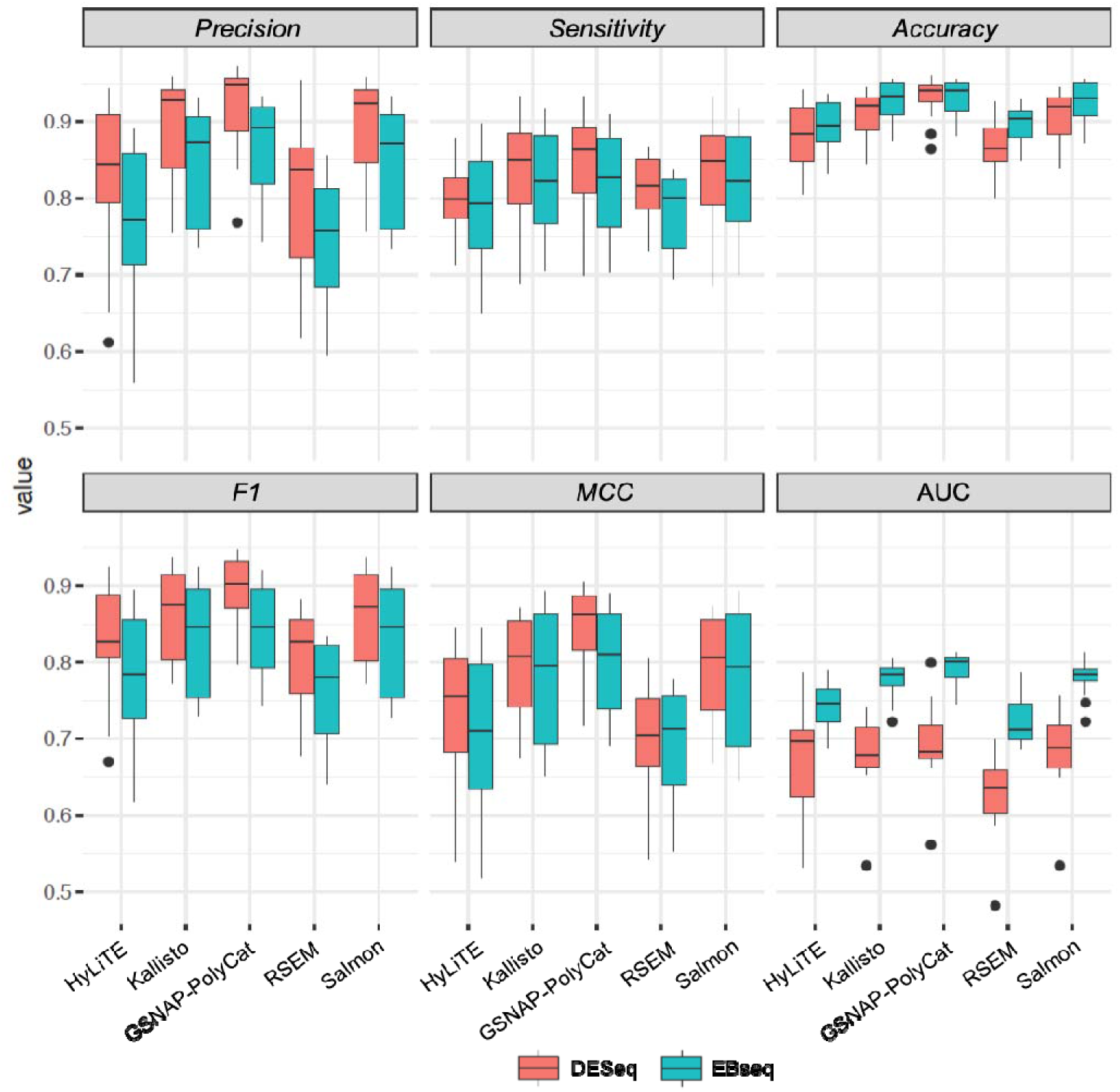
Performance evaluation of differential expression analysis between homoeologous genes.

Co-expression relationships between homoeologs were measured using Pearson’s correlation coefficient across multiple sample conditions. Approximately 1-5% of homoeolog pairs (418-1,834 out of 37,223 pairs) exhibited significant changes due to incorrect read assignment. As shown in Figure 4, GSNAP_PolyCat and HyLiTE introduced the smallest numbers of differential co-expression (DC) changes, thereby outperforming RSEM, Salmon, and Kallisto. Artifactually-induced DC was most prominent in those homoeologous gene pairs exhibiting higher *Ambiguity* (Figure 2G), with the highest bin (i.e., 0.2-1) exhibiting a nearly 4-fold increase in DC than other bins for RSEM, Salmon, and Kallisto. Among the nine categories of DC changes (Figure 4, columns), the class of 0/+ was most significantly enriched for each pipeline except for GSNAP-PolyCat, where it was the second most enriched category after +/0. This suggests that the majority of DC changes due to read partitioning errors lead to gains in correlation, generally changing our inferences from no significant correlations (0) to significantly positive correlations (+). These observations suggest read partitioning methods could lead to an over-estimation of co-regulation between homoeologous genes due to incorrect homoeolog expression estimation, consequently restricting our ability to infer expression divergence and/or possible functional divergence of duplicated genes. Notably, these patterns were consistent for both the *rlog* and log_2_RPKM data transformation methods. In addition to DC between homoeologous gene pairs, we also conducted identification and classification of DC patterns for all possible gene pairs (Supplementary Table S3), resulting in 0.3%-1.1% global pairwise DC changes, which affected 9.3-15.2% of total genes (i.e., DC genes enriched with DC pairs) in their co-expression relationships.

**Figure 4.**
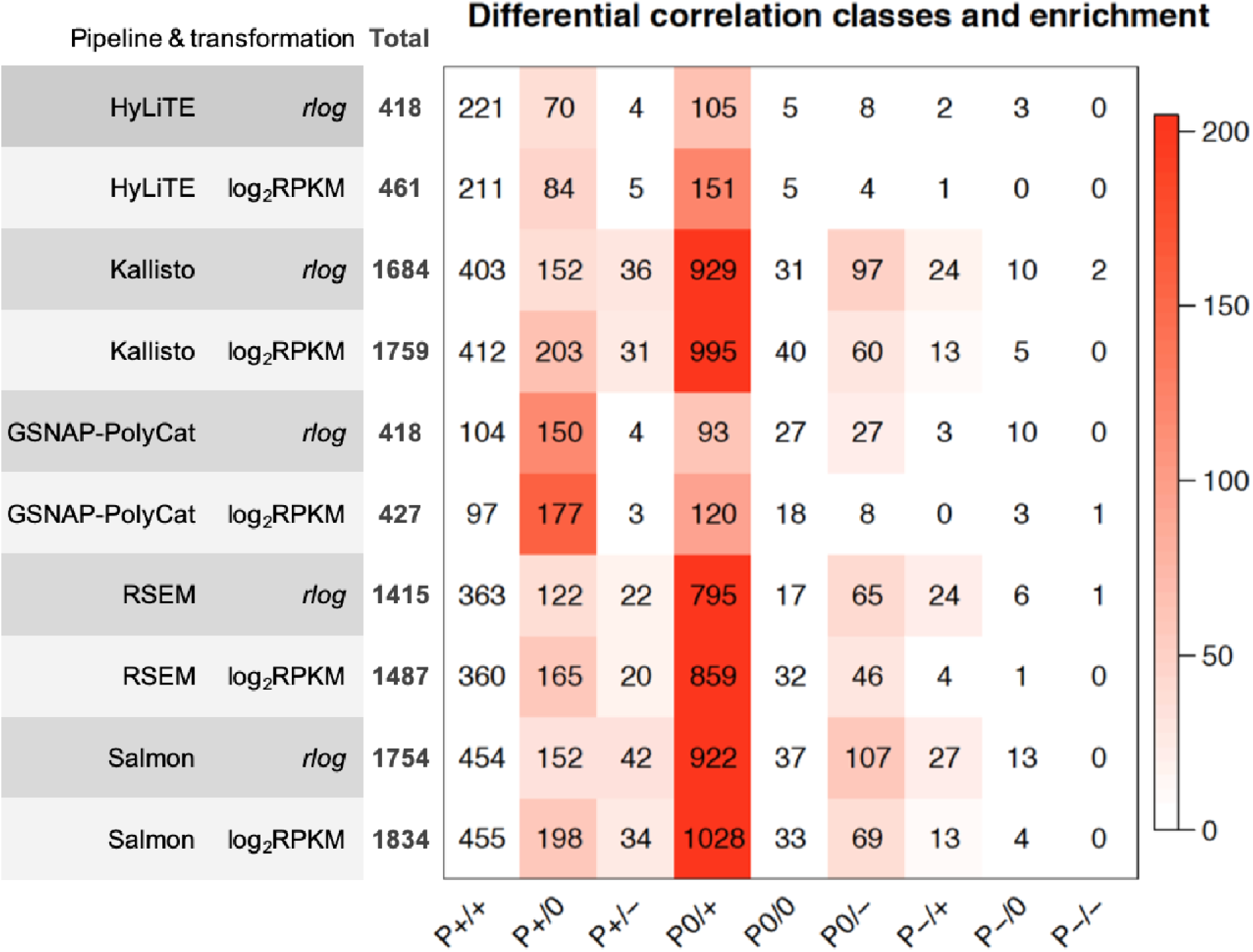
Inference changes in co-expression relationships between homoeologs. For each of the 10 combinations of homoeolog expression estimation pipelines and data transformation methods (row), the number of differential correlation (DC) changes between observed and expected datasets are shown for each DC category (column). Cell color indicates the magnitude of significant over-representation based on -log_10_(*P*-value) of Fisher’s exact test (i.e., *P* = 0.05 is converted to 1.3). For example, the number in category P0/+ of the bottom row indicates that 1028 homoeolog pairs showed DC changes from no significant correlation (0) to significantly positive correlation (+) due to the estimation error from the Salmon pipeline followed by log_2_RPKM transformation.

### Robust construction of gene co-expression networks by the rank-based binary method and WGCNA

Gene co-expression networks are commonly used to summarize the multidimensionality of gene expression data into clusters of genes with putatively related functions (i.e., modules). In the context of polyploidy, co-expression networks can be used to assess functional relatedness among genes and homoeologs, reveal changes in homoeolog usage, and characterize the genetic interplay between subgenomes [6]. We use both weighted and unweighted networks to assess the influence of variation in read partitioning on our inferences of coexpression.

Constructing un-weighted co-expression networks requires a binary classifier (or hard threshold) to decide whether there exists a connection (i.e., an edge) between each pair of genes. As shown in Figure 5A, different rank-based thresholds (5%, 1%, 0.5% or 0.1% of top ranked correlations become edges) yielded robust classification of the expected edges (based on diploid expression) with AUC scores close to 1. In contrast, the performance of *Z*-statistics-based thresholds (i.e., significant correlations with *Z*-score above 1.5, 2.0, 2.5, or 3.0 become edges) were more variable (AUC of 0.8∼1) depending on the stringency of the Z thresholds. These results indicated that the rank-based method is more robust here than *Z-statistics* to infer binary gene co-expression network.

**Figure 5.**
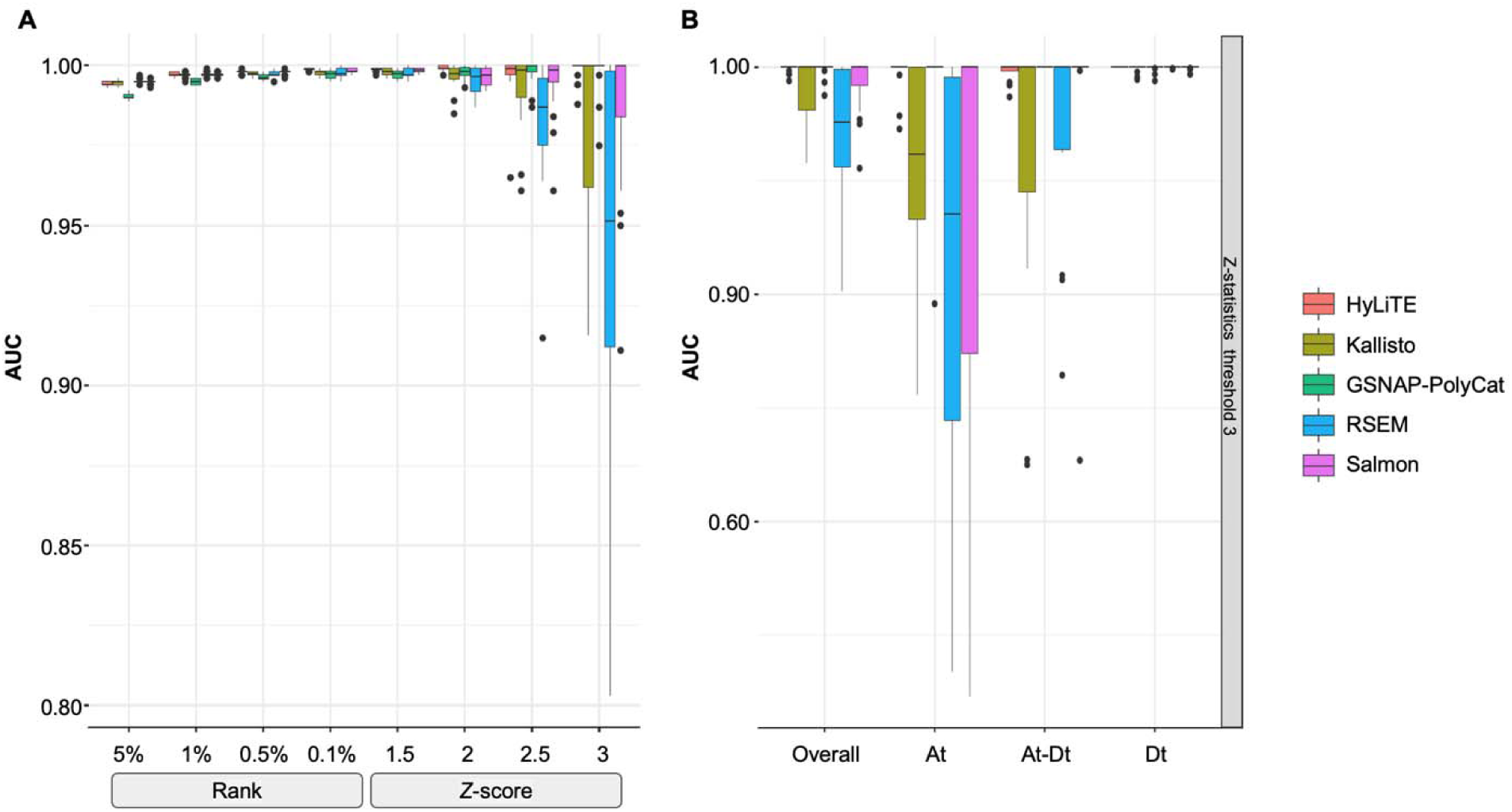
Performance of binary co-expression network construction. **A**. Boxplot of AUC scores were shown using different homoeolog estimation pipelines (color) and binary thresholds (x-axis). **B**. Taking the *Z*-score threshold of 3, for example, AUC scores were compared among subnetworks: Overall - all edges considered; At - edges within the A-genome subnetwork; Dt - edges within the D-genome subnetwork; At-Dt - edges connecting genes across A- and D-subnetworks.

In addition to the network construction methods (ANOVA formula: AUC ∼ construction + pipeline + transformation; construction *P* < 2e-16), the choice of read estimation pipeline also matters (*P* = 3.91e-09) with performance of RSEM significantly falling behind others (Tukey’s HSD test *p* < 0.05); no significant difference was found between the *rlog* and log_2_RPKM transformation (ANOVA = 0.624). Interestingly, while not unexpected, edge inference within the D-genome subnetwork is significantly more robust than edges within the A-genome subnetwork or those across subnetworks (Figure 5B; ANOVA and Tukey’s HSD test < 0.05). This observation likely reflects quality differences in the mapping reference, i.e., the high quality D-genome reference and the inferred A-genome sequences (see methods).

In weighted gene co-expression analysis (WGCNA) networks, the quantitative strength of network connections is considered to maximize information captured in the network. The topological preservation tests of expected modules (based on diploid expression) exhibited high preservation (*Z_summary_* > 10) for almost all modules (Supplementary Figure S1A), regardless of soft threshold (i.e., 1, 12, or 24; see methods), homoeolog read estimation pipeline, and method of normalization. This result suggests that WGCNA-based inference of gene modular structure is rather robust.

In addition to the separate topological evaluation above (binary networks by edge inference AUC and WGCNA networks by module preservation), node connectivity (*k*) and network functional connectivity (*FC*) were calculated for each binary and weighted co-expression network constructed. Each AD network constructed was evaluated against the expected (diploid-based) network. Pearson’s correlation coefficients between the expected and observed networks suggest that both *k* and *FC* were rather consistent across different homoeolog expression estimation pipelines (ANOVA formula: correlation ∼ construction + pipeline + transformation; construction *p* > 0.05), whereas the method of network construction could strongly influence topology (*p* < 2e-16; Supplementary Figure S1B-D). Notably, normalization method affected *k* (*p* < 2e-16; log2RPKM outperforms *rlog*) but not *FC* (*p* = 0.08). As shown in Supplementary Figure S1B-D, both rank-based binary construction and weighted gene network construction methods equally outperformed all but the least strict *Z*-statistics methods. The accurate inference of *k* (measured by correlation between observed and expected data; Figure 2H) is negatively correlated with *Ambiguity*, albeit weakly. This diminished relationship is expected as the network property of each gene is intrinsically determined by all the other genes, thereby obscuring the impact of ambiguity per gene.

In addition, the measure of *FC* can be used to statistically evaluate the functional significance of network topology [54]. According to the “Guilt-by-Association” principle [59], genes with similar functional properties tend to interact or be clustered together in biological networks. Thus, higher *FC* indicates more reasonable network topology. As shown in Supplementary Figure S2, the highest *FC* scores were observed for WGCNA networks (AUROC *=* 0.63-0.66), followed by the ranked-based binary networks (0.54-0.63) and the *Z*-statistics-based binary networks (0.50-0.54), respectively. This may suggest that the WGCNA network construction was able to capture more function and/or biologically-relevant information.

Overall, the performance of co-expression network analysis was more affected by network construction methods than by read ambiguity and partitioning methods. In general, either log_2_RPKM or *rlog* combined with WGCNA produced the best results for these data, regardless of read assignment method.

### Bioinformatic choices can strongly affect the interpretation of duplicated gene network topology

In the context of polyploid gene network, it is of particular interest to compare subnetwork properties within each subgenome and between subgenomes. Taking the GSNAP-PolyCat dataset followed by log_2_RPKM normalization as an example, both rank-based and WGCNA networks (Figure 6A and 6C, respectively) revealed the highest density (mean connections) of the A-subnetwork, followed by that of the D-subnetwork and then of the inter-connections between A and D subgenomes. In contrast, similar levels of A- and D-subnetwork density were revealed in the *Z*-statistics-based networks (Figure 6B). These results led to opposite conclusions regarding the potential topological asymmetry between two subgenomes. According to the performance assessment above, we believe that the conclusion derived from WGCNA and rank-based binary networks is more reliable; that is, the At genes are more interconnected than are the Dt genes, reflecting the difference in gene regulation between the two subgenomes (i.e. the A_2_ and D_5_ diploids used generate synthetic AD). In addition, all networks agreed on the much lower density of inter-subgenome connections than that of within-subgenome connections, indicating that a gene is much more likely to be connected with genes from the same subgenome than with genes from the other subgenome. For other combinations of homoeolog expression estimation, transformation and network construction methods, the measures of subnetwork density are shown in Supplementary Table S4.

**Figure 6.**
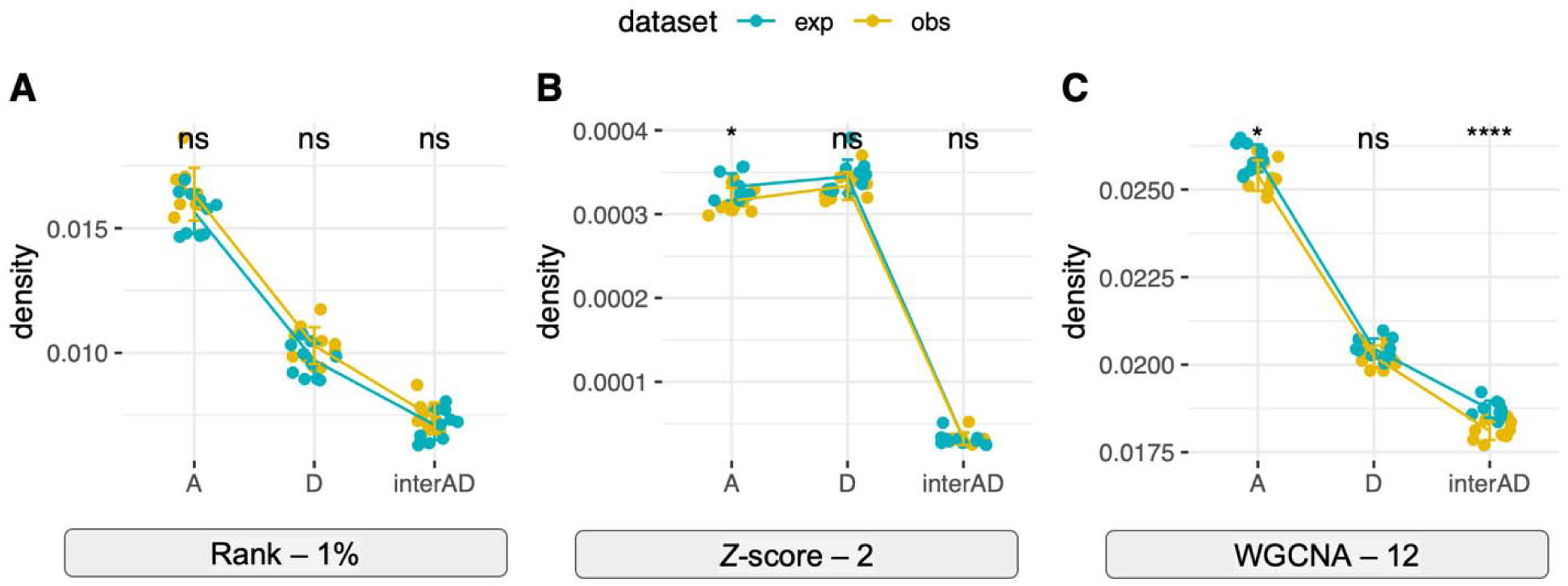
Different inferences of subnetwork topology. The network density of A-subnetwork, D-subnetwork, and interconnections between A- and D-subnetworks were shown for both the expected and observed data from the GSNAP-PolyCat estimation with log_2_RPKM normalization. **A** - rank-based binary network with top 1% connections; **B** - *Z*-statistics binary network with connections above *Z*-score of 2; **C** - WGCNA network with power of 12.

## DISCUSSION

The duplicated nature of polyploid genomes poses unique challenges for bioinformatics. Presently, we are witnessing an explosion of interest in better understanding these challenges and developing appropriate methodologies and tools for polyploids, for applications as diverse as genome sequence assembly [60], genotyping [61, 62], haplotype phasing [63, 64], population-based trait analysis [65], phylogenetic inference [66, 67], and transcriptomic-based analyses [68, 69] such as *de novo* transcriptome assembly [70] and transcript quantification [71]. Quantification of homoeolog expression is particularly interesting, given the various patterns of duplicate gene expression possible in polyploid species (reviewed in [12]), the interactions among homoeologs in a gene network context [6, 14], and the phenomenon of unbalanced homoeolog expression bias together with its potential long-term consequences for fractionation [72–74]. A number of previous studies have explored the bioinformatics of homoeolog expression profiling [68–71]; however, both the fundamental issue of read ambiguity and the downstream inferences regarding polyploid expression evolution have not been addressed. Here we present a comprehensive analytic workflow to demonstrate the challenges and pitfalls of these analyses (Figure 1), as well as how they are influenced by the extent of read ambiguity in the dataset and how that ambiguity is handled in understanding homoeolog expression and co-expression patterns (Figure 2).

### Duplication and deficiency: when redundancy renders reads unresolved

In addition to the redundant nature of polyploid genomes, there are a number of biological and technical causes for ambiguous read mapping, including transcripts that are expressed at low levels, sequence homology, small-scale gene duplications, and errors in sequencing and annotation. While we can control some of these factors through experimental design (i.e., read length, paired-end sequencing, etc), the nature of the biological system and the amount/distribution of subgenome divergence, as measured by *Ambiguity*, will influence the ability to accurately assign reads to homoeologs. Although our analysis is limited to the example data from *Gossypium*, the metric of *Ambiguity* can be applied to any other real-world or simulated polyploid systems. For systems that have less divergent subgenomes than *Gossypium*, the *Ambiguity* values are expected to be higher, and longer read lengths will be required to improve the ability to accurately assign reads. Knowing the range of *Ambiguity* for any specific polyploid system or for a list of genes of interest, we can foresee the use of Figure 2 to query how such a range) affects the performance of bioinformatic inferences regarding homoeolog read estimation (B-E) and polyploid expression evolution (F-H).

Among tools that have been devised to estimate homoeolog expression levels under different conditions (e.g. the availability and type of the reference genome; Figure 7), numerous methods exist for handling the subset of reads that are not uniquely assignable, typically either discarding these reads (as in GSNAP-PolyCat and HyLiTE) or statistically assigning the reads (e.g., RSEM, Kallisto, and Salmon). Among the five pipelines evaluated in this study, most performed relatively well, achieving >90% success for information retrieval metrics. Notably, GSNAP-PolyCat exhibited the best scores for most metrics, aside from those affected by read removal (i.e., *Efficiency* and *Discrepancy*). While it is tempting to attribute the improved performance of this pipeline to the underlying resequencing-based SNP data, which was not used by HyLiTE, the remaining pipelines (i.e., RSEM, Salmon, and Kallisto) were all provided a reference transcriptome derived from the homoeoSNP index used in GSNAP-PolyCat. When *Ambiguity* was low, all pipelines performed similarly well; however, those that statistically assign ambiguous reads (RSEM, Salmon, Kallisto) perform significantly worse for those genes with *Ambiguity* above 20%. This may be due to the noise in the underlying statistics as the relative number of unique reads drops compared to those that will be statistically assigned; that is, any error in statistical inference will be amplified as the number of ambiguous reads begins to outweigh the number of unique reads. This is an important observation for polyploid systems whose subgenomes are more recently diverged. That is, methods which statistically assign ambiguous reads should be used with caution when the divergence between parental genomes is low. For those genomes, GSNAP-PolyCat and HyLiTE will provide a more reliable representation of relative homoeolog read counts, with GSNAP-PolyCat outperforming HyLiTE when *a priori* homoeoSNP information is available.

**Figure 7.**
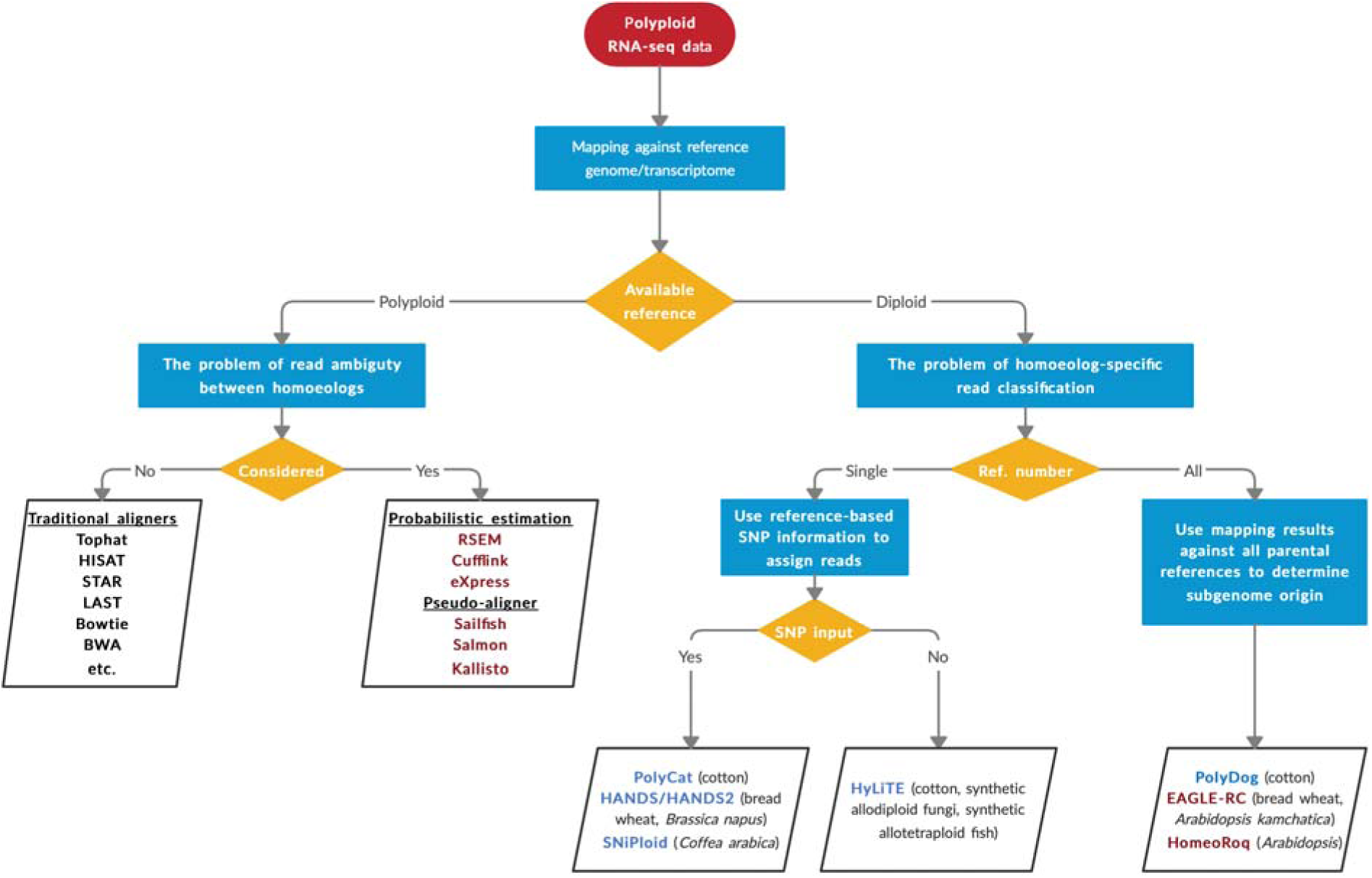
A decision-making diagram to choose appropriate bioinformatic resources for estimating homoeolog expression levels. When a reference genome or transcriptome is available for the polyploid species, quantification of individual homoeologs is either straight-forward using the traditional aligners such as Tophat, or applying probabilistic estimation methods and pseudo-aligners to consider the problem of read ambiguity. When the reference is only available for one or more diploid progenitors, software has been developed for partitioning and/or quantifying homoeolog-specific reads: maroon colored software, such as RSEM and EAGLE-RC, statistically assign the subgenome origin for ambiguously mapped reads; blue colored software such as PolyCat utilize only unambiguously mapped reads for estimation. The polyploid systems for which they were originally developed are noted in parentheses.

In a previous study, [71] showed that EAGLE-RC, a likelihood model-based method, outperforms other homoeolog expression quantification methods including STAR, LAST, Kallisto, and HomoeoRoq to precisely estimate homoeolog expression in both tetraploid *Arabidopsis kamchatica* and hexaploid wheat. The category of the subgenome-classification approaches (Figure 7, bottom right), including EAGLE-RC, HomeoRoq [75], and PolyDog [22], requires read mapping against each subgenome separately in order to determine the better supported homoeolog origin for reads. These approaches were not included in our study, because the reference quality and annotation methods differ between the A- and D-diploid progenitor genomes, which introduces additional dimensions of complexity for homoeolog quantification. For example, GSNAP-PolyCat and HyLiTE appeared to partition more reads than expected to the higher quality D-genome reference, whereas the other three pipelines statistically characterized more reads as A-derived; the cause (likely differences in algorithm) of this discrepancy is unknown, but it has consequences even at the co-expression network level (more robust inference of D- vs. A-subnetwork topology). These caveats notwithstanding, we envision that this category of approaches will be useful for hybrid and polyploid systems where quality differences among progenitor reference genomes are negligible and where similar annotation methods are used for each.

### Consequences of inaccurate quantification for inferences of polyploid evolution

Beyond the narrow issue of evaluation of homoeolog quantification, our interest lies in identifying a reasonable set of methods to address biological and evolutionary questions concerning polyploidy. Differential expression (DE) is commonly among the first transcriptomic analyses performed, providing a generalized look at the extent of expression divergence. For polyploid species, the relative expression of each subgenome is of particular interest, which may provide insights into homoeolog bias, expression level dominance, *cis*-*trans* resolution, putative sub-/neo-functionalization of homoeologs, and other phenomena [14]. In general, GSNAP-PolyCat best represented the expected DE between homoeologs, followed closely by Kallisto and Salmon (Figure 3). The standout, RSEM, performed significantly more poorly than the rest despite its intended application to isoform quantification; we therefore advise caution when using RSEM for duplicate gene analyses. With respect to DE inference, both DESeq2 and EBseq resulted in reasonable performance metrics, with the choice likely being the stringency level and parameters preferred.

In contrast to the general robustness of the DE results, homoeolog read ambiguity and the choice of quantification pipelines strongly influence our interpretation of co-expression relationships among genes. In particular, we are interested in detecting coordination among homoeologous genes in polyploids. The most significant error evident is the false detection of positive correlations where none exist. Notably, those methods that discard reads (i.e., GSNAP-PolyCat and HyLiTE) far outperformed the other methods, particularly for those genes with higher *Ambiguity*. These results were consistent for the two normalization methods tried, i.e., *rlog* and log2RPKM, and may indicate a general preference for discarding ambiguous reads when the biological question depends on an accurate assessment of differential co-expression.

The inference of co-expression network topology, on the other hand, was generally less sensitive to the quantification method, but rather was dependent on method of network construction. This is probably because the multivariate nature of co-expression relationships mitigates the influence of individual and random quantification errors. In order to compare network topologies between subgenomes, choosing the appropriate network construction method becomes critical, otherwise incorrect and even opposite conclusions may be reached (Figure 6). For example, both rank-based binary and WGCNA reconstructions of the present datasets suggest that the A-subnetwork is more tightly interconnected than the D-subnetwork, whereas the less reliable *Z*-statistic based binary networks suggest they are equally interconnected.

## Conclusions

In this study, we present an analytical workflow from homoeolog expression quantification to a series of downstream analysis to infer key phenomena of polyploid expression evolution. By examining the extent and consequences of read ambiguity, we demonstrated the potential artifacts that may affect our understanding of duplicate gene expression, such as an over-estimation of homoeolog co-regulation and the incorrect inference of subgenome asymmetry in network topology. Such errors may be reduced by mitigating technical factors that influence ambiguity, i.e. sequencing strategy and fundamental resources (i.e., genomes and/or resequencing). Although the collection of methods tested in this study may be superseded by those yet to be developed, our work introduces the metric of *Ambiguity* and designates a set of reasonable practices applicable to other polyploid systems.

## KEY POINTS

- We present an analytical workflow to evaluate a variety of bioinformatic method choices at different stages of polyploid RNA-seq analysis, from homoeolog expression quantification to downstream analysis used to infer key phenomena of polyploid expression evolution.
- We used transcriptomic data from the cotton genus (*Gossypium*) as an example to examine the extent and consequences of homoeolog read ambiguity.
- Our results show that GSNAP-PolyCat outperforms other quantification pipelines tested, and its derived expression dataset best represents the expected results in downstream analyses of differential expression and co-expression network analysis.
- We illuminate the potential artifacts that may affect our understanding of duplicate gene expression, including an over-estimation of homoeolog co-regulation and the incorrect inference of subgenome asymmetry in network topology.
- Overall, our work points to a set of reasonable practices that are broadly applicable to the evolutionary exploration of polyploids.

## ACKNOWLEDGEMENTS

We thank the National Science Foundation Plant Genome Research Program and Cotton Incorporated for financial support. We also thank ResearchIT for computational support at Iowa State University.

## SUPPLEMENTARY DATA

**Supplementary Figure S1.** Evaluation of module preservation, node connectivity and functional connectivity (GO and KEGG) between observed and expected polyploid networks. For module preservation tests of WGCNA networks (upper left), *Z_summary_* > 10 indicates strong preservation.

**Supplementary Figure S2.** Comparison of network functional connectivity across homoeolog expression estimation pipelines and co-expression network construction methods.

**Supplementary Table S1.** Summary of the cotton RNA-seq datasets in this study.

**Supplementary Table S2.** Observed and expected expression divergence between At and Dt homoeologs.

**Supplementary Table S3.** Summary of differential co-expression analysis.

**Supplementary Table S4.** Measures of subnetwork density.

