## Supplementary figures and images for "Homoeologous gene expression and co-expression network analyses and evolutionary inference in allopolyploids"

### Figure S1

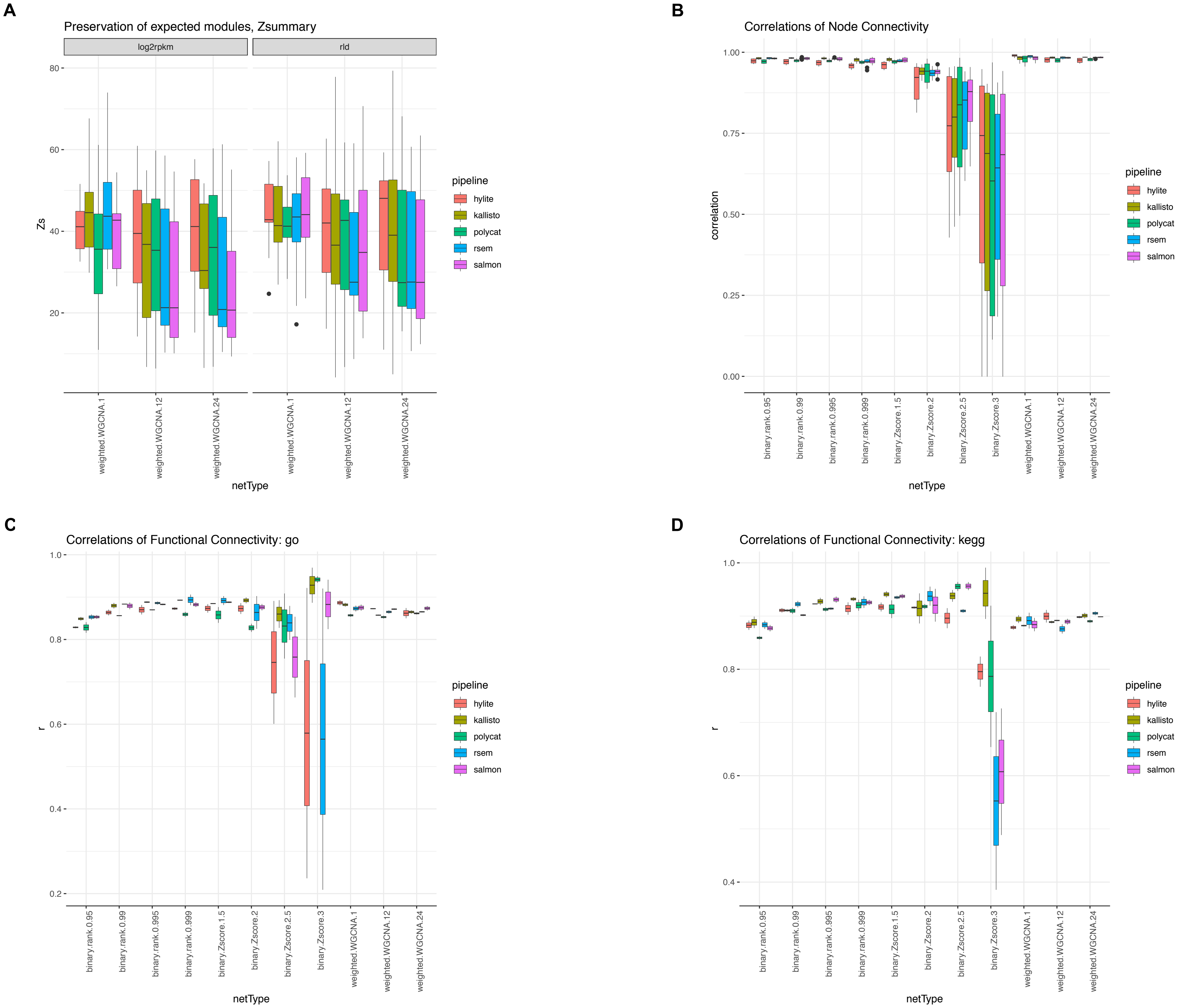

### Figure S2

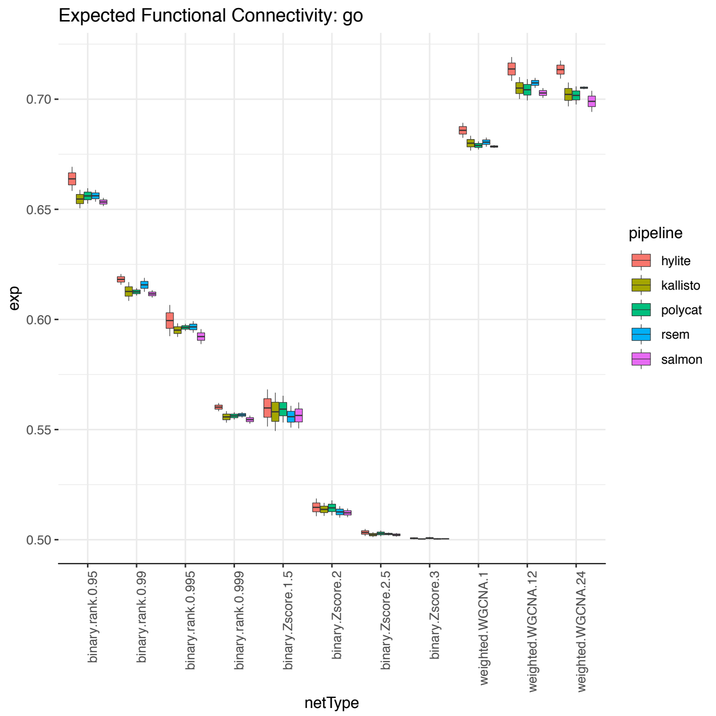

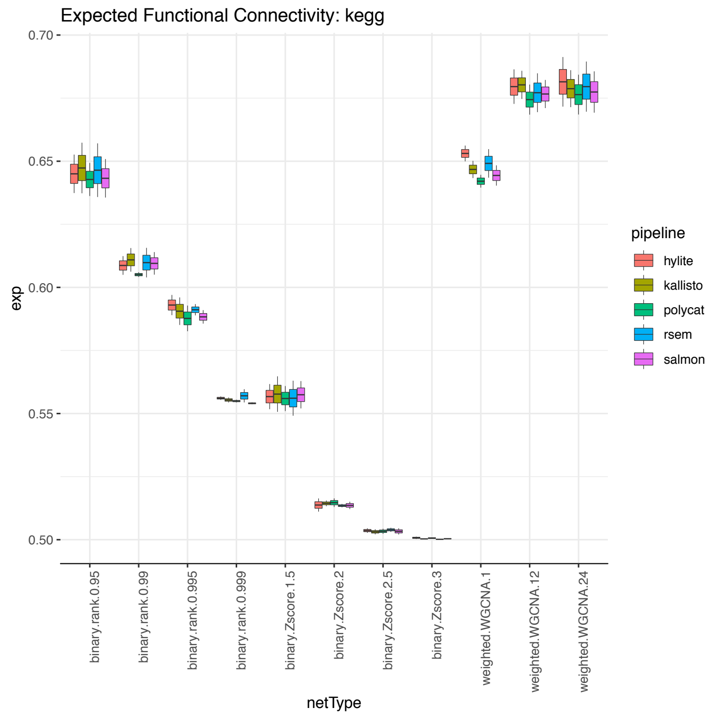

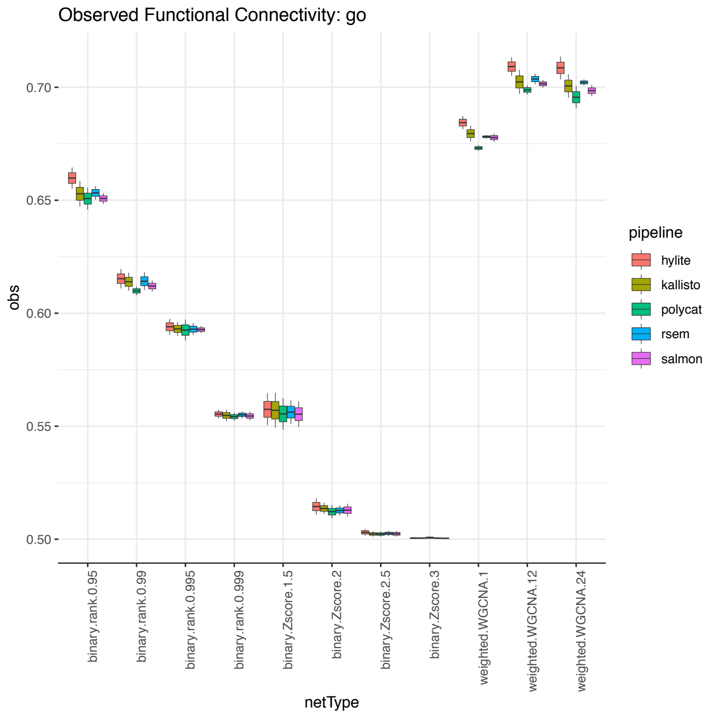

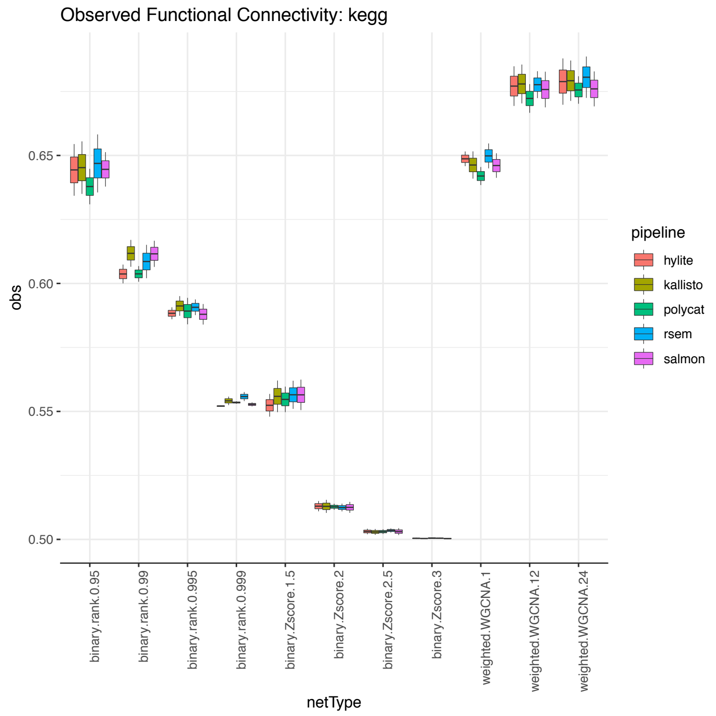
